# 3D flightpaths reveal the development of spatial memory in wild hummingbirds

**DOI:** 10.1101/2022.05.18.492296

**Authors:** David J. Pritchard, T. Andrew Hurly, Theoni Photopoulou, Susan D. Healy

## Abstract

Many animals learn to relocate important places and reflect this spatial knowledge in their behaviour. Traditionally evidence for learning is examined experimentally by studying spatial memory. However, tools developed for analysing tracking data from widely ranging animals allow a more holistic analysis of behaviour. Here we use the two together in novel combination of experimental and modelling approaches to analyse how patterns of hummingbird movements change as birds learn to find a reward in a location indicated by a pair of landmarks. Using hidden Markov models (HMMs) we identified two movement states which we interpret as Search and Travel and compared these to experimental behavioural measures of spatial memory. When birds had a single training trial to learn a flower’s location, both the behavioural measures and HMMs showed that hummingbirds relied on landmarks to guide search. Hummingbirds focussed hovering around the rewarded location and were more likely to be in the Search state, and more likely to switch from Travel to Search, when closer to the rewarded location, but only when the landmarks were present. When birds had had 12 additional training trials, however, the HMMs and behavioural measures showed differences in how reliant birds were on landmarks. While behaviours like hovering were still strongly affected by removing landmarks, the likelihood of being in or entered the Search state was the same regardless of whether the landmarks were present or removed. These results suggests that hummingbirds rapidly learn to use nearby landmarks to structure where they search, but as birds gain experience the role of these landmarks changes. While familiar local landmarks were still essential for precise search, experienced birds were able to use alternative cues to guide broad-scale transitions between behaviour. HMMs and traditional behavioural measures each capture a different aspect of this learning, with neither approach alone accurately described the role of landmarks in spatial learning.

## RESULTS AND DISCUSSION

For many animals spatial memory enables accurate return to rewarding foraging locations^1^. Flower- visiting species also use spatial memory to avoid flowers they have recently emptied, which will not yet have replenished their nectar supplies^2^. Experimental manipulations have supplied most of our understanding of how animals acquire and use spatial memory: animals are trained to visit a particular, rewarding location and researchers measure where animals focus their search once the reward is removed (e.g. where animals choose to dig^3,4^ or peck^5^). These measures focus on the endpoints of search because the endpoints can be easily quantified and converted into measures of accuracy (e.g. proximity to the goal). Comparisons of changes in accuracy over multiple trials can reveal how spatial learning develops^3,6^. Tracking studies of animals face the opposite problem. Continuous tracking of animal movements provides a rich insight into how behaviour changes over time, and ecologists have developed analyses to examine how movements change from moment to moment, examining both the spatial and temporal aspects of searching^7,8^. But while experimental studies can train animals to expect a reward in a particular location, providing strong a priori predictions about where animals using spatial memory should search, most tracking studies are purely observational and so discriminating between search based on memory and search based on real-time sensory information is challenging^9^.

Here we bring these two approaches together to identify how spatial memory shapes search. We do this for two reasons: first, while decades of experiments on spatial memory have shown how discrete choices become more accurate over time, little is still known about how spatial memory affects decisions before the animal reaches the endpoint. Analyses developed for studying animal tracks, on the other hand, provide an ideal toolkit for examining how spatial memory influences a much wider range of movements. Second, while movement ecologists have become increasingly interested in how learning and memory could shape animal movement^9,10^, here we validate these ideas by examining movement in a controlled environment thereby ruling out confounding cues and testing predictions about where animals should search.

To bring these two approaches together, and to identify how spatial memory shapes search, we take advantage of hidden Markov models (HMMs), which have been used by movement ecologists to identify changes in animal movement^11,12^. These tools classify animal paths into different “movements states”, based on properties such as speed and tortuosity. Such classification allows us to model how environmental factors influence the likelihood that the animal is in a state or is transitioning from one state to another. Although developed to examine the tracks of remotely- observed animals, HMMs are well suited to analysing how behaviour changes in experimental settings. In an experimental setting, by training animals to expect food in a particular location we can predict where animals should search if they use spatial memory. The trained animal’s behaviour also allows confidence that their behaviour is actually “search” rather than other superficially similar behaviours such as navigating difficult terrain or interacting with other animals.

To do this we tested how wild rufous hummingbirds *Selasphorus rufus* modify their behaviour as they learn about a location. Using a combination of HMMs and measures of search accuracy, we identified behavioural signatures of spatial memory in the 3D flightpaths of the hummingbirds, including how spatial memory shaped the endpoints of search, as well as where and when birds switched between different movement states. By manipulating the birds’ experience of a location (one experience or multiple experiences), as well as whether they had access to familiar landmarks, we experimentally tested how visual landmarks shaped how hummingbirds search. As a result we could see how these developed from rapid changes in movement to precise search as the birds gained more experience of a location.

### Landmark memory shapes movement after a single experience

To look at the effect of visual landmarks on search movements after a single experience, we trained territorial hummingbirds to visit an artificial flower filled with 25% sucrose solution and 30cm away from a pair of landmarks. We allowed each bird a single visit to the flower. In the test, we removed the flower and allowed the bird to search around the landmarks for the missing flower. All birds in Experiment 1 received the same test.

In Experiment 2, the landmarks and flower were moved to a new location and birds again had a single training trial to learn the flower’s location. In the test trial, for half the birds we removed the flower but not the landmarks (“landmarks present”). For the remaining birds, however, we removed both the flower and the landmarks (“landmarks removed”).

We analysed the 3D paths of the birds using two approaches: 1) HMMs analysed the paths in their entirety, identifying 2 likely states, and computing how these changed with distance to the flower; and 2) specific behavioural data, including where birds stopped and hovered in place (“stops”) and how close birds flew to the flower’s location, were analysed using generalised linear models with landmarks group and experiment as covariates. As all birds received the same treatment in Experiment 1, this experiment provided a baseline to which we could compare behaviour in the Experiment 2 in which half of the hummingbirds had both the flower and the landmarks removed.

When the landmarks were present (both groups in Experiment 1 and the “landmarks present” group in Experiment 2), the birds’ behaviour in the test depended on their proximity to the missing flower’s location (Table 1, Figures 1-3). In Experiment 1, birds in both groups flew and stopped close to the flower’s location, with the location of the closest stops often being the closest that the birds flew to the flower’s location in the test. This “tuning” of behaviour around the flower’s location was also seen in the HMM results, as birds switched from a faster and more direct style of movement, hereafter referred to as “Travel” to a slower, more sinuous style of movements we refer to as “Search”, when they approached the flower’s location. The inverse was seen as birds moved away from the flower’s location, increasingly switching from Search to Travel when more than 1m away (Figure 1). Together, stopping behaviour and HMM-identified states provide compelling evidence that hummingbirds focussed their search around a rewarded location after a single experience.

**Table 1:** Model comparison results for HMMs fitted to hummingbird data. Model in bold is the best performing model according to AIC weight, whereas models with an AIC weight of less than 0.0001 have been removed but can be found in supplementary materials.

| Exp. | Model | Covariate(s) | On | Angle mean estimates | AIC | Weight |
| --- | --- | --- | --- | --- | --- | --- |
| **1** | **1.1** | **Flower distance** | **Transition probabilities** | **None** | **3922** | **0.947** |
|  | 1.2 | None | - | Pitch & Yaw | 3928 | 0.052 |
| **2** | **2.1** | **Flower distance x Landmarks** | **Transition probabilities** | **None** | **1723** | **0.973** |
|  | 2.2 | Flower distance, Landmarks | Transition probabilities (distance), mean step length (landmarks) | None | 1732 | 0.014 |
|  | 2.3 | Flower distance + Landmarks | Transition probabilities | None | 1734 | 0.004 |
|  | 2.3 | Flower distance | Transitions from traveling | None | 1734 | 0.003 |
| **3** | **3.1** | **Flower distance, Landmarks** | **Transition probabilities (distance), mean step length (landmarks)** | **None** | **1440** | **0.711** |
|  | 3.2 | Flower distance x Landmarks | Transitions from traveling | None | 1443 | 0.131 |
|  | 3.3 | Flower distance, Landmarks | Transitions from traveling (distance), mean step length (landmarks) | None | 1443 | 0.124 |
|  | 3.4 | Landmarks | Mean step length | None | 1445 | 0.003 |

**Figure 1:**
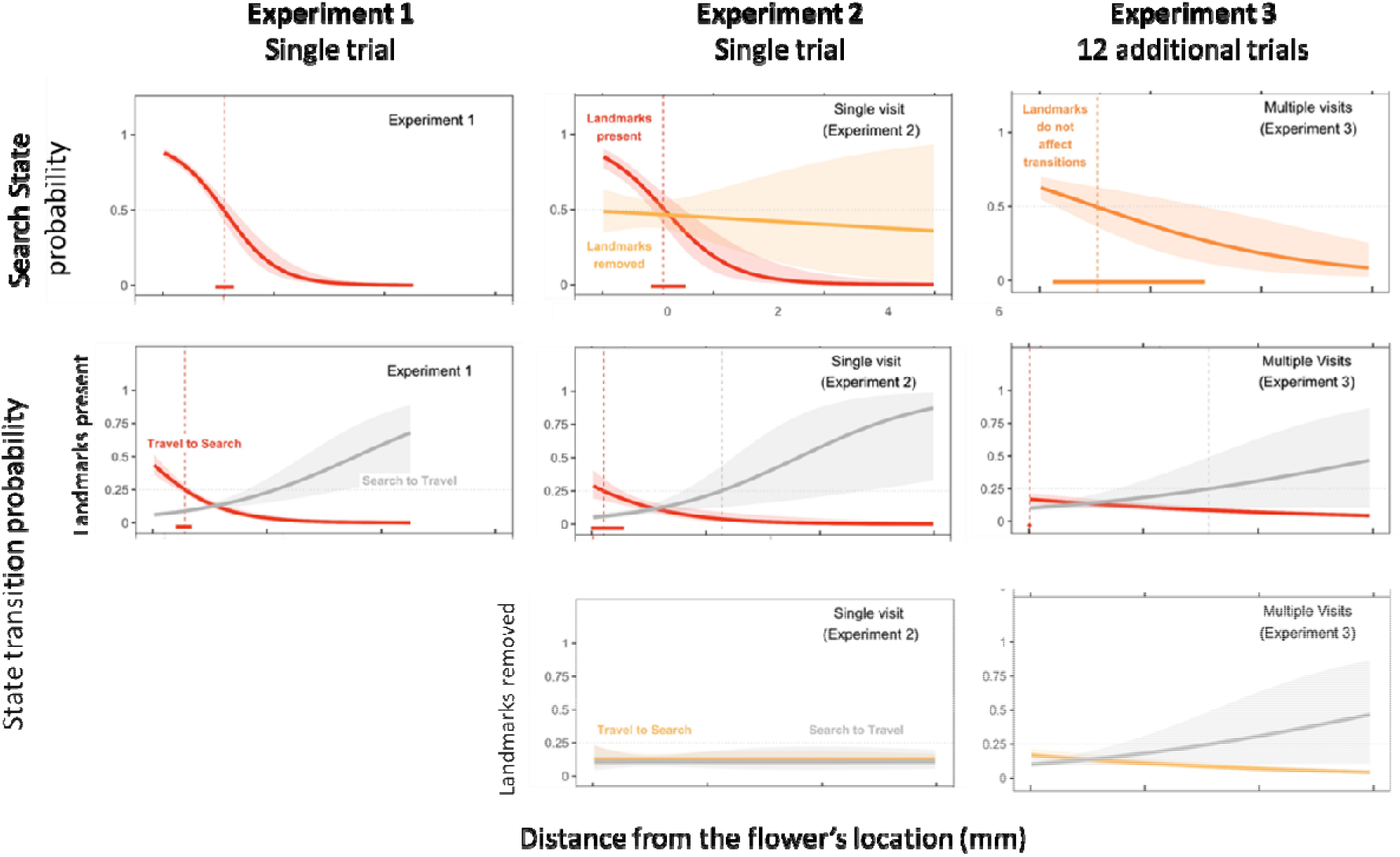
Hidden Markov Model results for how stationary state probability and transition probability changes with distance to the flower’s location. Top row: the probability of birds being in the Search State at different distances from the flower, when the landmarks are present (red line) or removed (yellow line). The dashed line and bar along the x-axis displays the distance (±95% confidence interval) where birds crossed over 50% probability of being in Search. In Experiment 3, no difference was found between landmark groups and so a single orange line represents both groups. Middle and bottom rows: he probability of birds transitioning between Search and Travel at different distances from the flower’s location when the landmarks are present (middle row) or removed (bottom row). The probability of ending search (Search -> Travel) is shown as a gray line while the probability of initiating search is represented as a coloured line (red for landmarks present, yellow for landmarks removed). The dashed line and bar along the x-axis displays the distance (±95% confidence interval) where birds crossed over 25% probability of being transition between their specified states.

This spatial tuning of behaviour to a rewarded location was, however, dependent on the landmarks. While the birds with the landmarks present in Experiments 2 behaved as they had in Experiment 1, birds with the landmarks removed drastically altered their behaviour. After the landmarks were removed, both the average distance of stops (t = -5.98, P < 0.0001) and the average distance during the entire test (z = -17.85, P < 0.0001) were further from the flower’s location than they had been in Experiment 1 (Figure 2). The distance of the closest stop (z = -4.39, P < 0.0001) and the closest distance a bird flew to the flower (z = 2.54, P = 0.011) were also further away, suggesting that the change in average distances represent a decrease in accuracy rather than an increase in variation.

**Figure 2:**
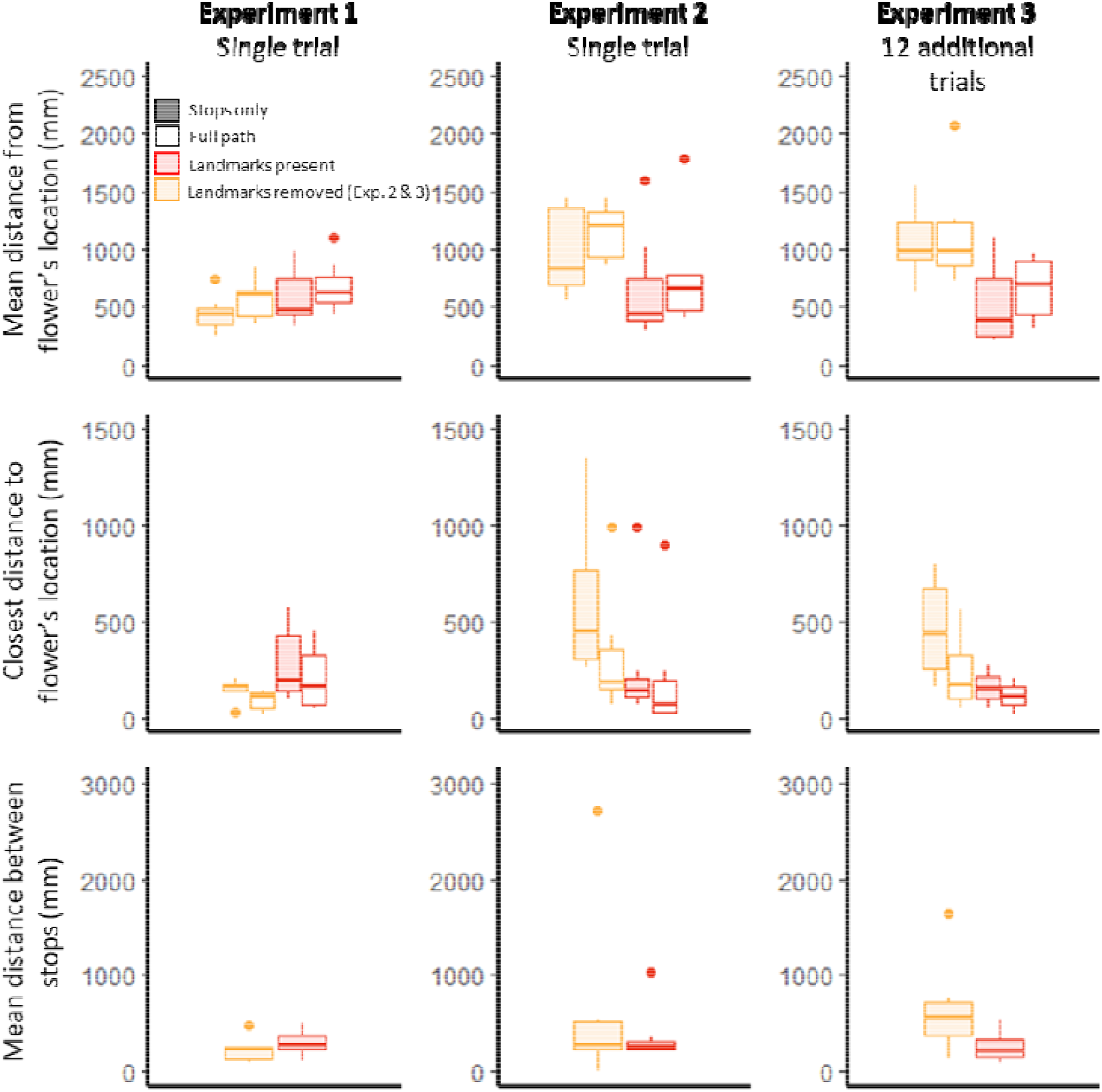
Behavioural measures of searching in the test. Top row: the mean distance that birds stopped (filled) and flew (open) to the flower’s location when the landmarks were present (red) or removed (yellow). Middle row: the closest distance that birds stopped (filled) or flew (open) to the flower’s location when the landmarks were present (red) or removed (yellow). Bottom row: the mean distance between adjacent stops when the landmarks were present (blue) or removed (red).

The birds without the landmarks were not just less accurate. They also changed how they stopped. The birds with the landmarks present in Experiment 2 spaced their stops relatively close together in both Experiment 1 and Experiment 2, with most stops within 30cm of the previous stop (landmarks present group, mean + se: Experiment 1: 305.8+-41.7mm; Experiment 2: 307.8+-53.7mm, Figure 2). The birds with the landmarks removed in Experiment 2, however, increased the distance between stops to by almost 20 cm, to almost 40cm between stops, between Experiment 1 and Experiment 2 (landmarks removed group, mean +- se: Experiment 1: 204.3+- 28.8mm; Experiment 2: 388.7+-88.3mm; experiment*landmark interaction: t = -2.36, P = 0.018). For this group the effect of distance from the flower disappeared as birds were equally likely to be searching or travelling at all distances from the flower’s location. Removing the landmarks therefore not only affected accuracy, but also resulted in changes in *how* birds searched.

### Experience affects movement states and stopping behaviour

To look at the effect of experience on hummingbird search, in Experiment 3, the birds were allowed 13 training trials before the test. All birds had landmarks present in the training trials but at test, as in Experiment 2, we removed the landmarks for one group of birds (the same birds as in Experiment 2).

As before we analysed how landmarks affected behaviour by using HMMs to examine how distance from the flower affected behavioural states, and by comparing birds’ behavioural measures between Experiment 2 and Experiment 3. As the landmark treatment (present versus removed) did not change between Experiments 2 and 3, we expected that any change we observed in behaviour was likely due to the birds having experienced 12 additional training trials.

We found that additional experience did indeed change how reliant birds were on the local landmarks: in Experiment 2 only the birds with the landmarks present were more likely to be in the Search State when closer to the flower, while in Experiment 3 both groups of birds (with and without the landmarks) increased the likelihood of Search when closer to the flower’s location. While there were slight differences in how fast experienced birds during the Travel state, in Experiment 3 the local landmarks were no longer necessary for the tuning of a bird’s Search State to the flower’s location (Figure 1, Table 1). With the additional training the birds had apparently learned to use features of the environment other than the landmarks we provided.

While experienced birds apparently no longer needed the local landmarks to enter the Search state, even with 12 additional trials the landmarks still significantly influenced how birds behaved when searching. The birds in Experiment 3 with the landmarks still tended to stop (landmark group: z = 1.74, P = 0.08) and tended fly (landmark group: z = 1.80, P = 0.07) on average closer to the flower’s location than did the birds without the landmarks (Figure 2). The distribution of stops also changed as the birds gained experience: while birds without the landmarks spaced their stops slightly further apart with experience, birds with the landmarks spaced theirs slightly closer together (z = 1.87, P = 0.062, Figure 2). Experienced birds, then, used the landmarks to focus their search for the flower’s location, and removing the landmarks resulted in birds both shifting their search further from the flower’s location and spreading search over a wider area, resulting in less focussed searching (Figure 3).

**Figure 3:**
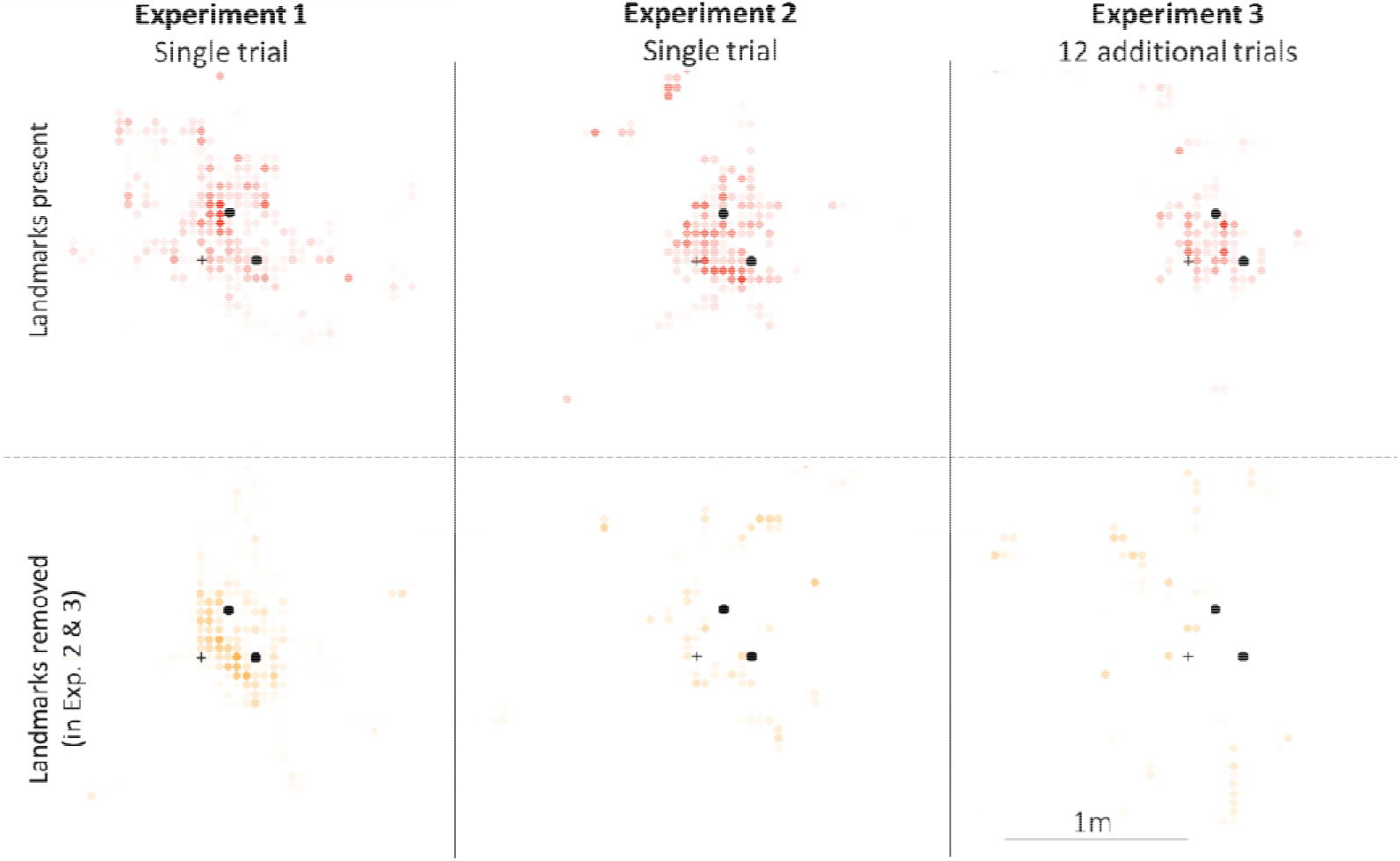
Heat-maps of the location of datapoints assigned to the Search state by the viterbi algorithm for the most successful model. Space was divided into 5cm diameter squares, the black dots are the landmarks and the flower’s location is represented as a +. Darker points mean that more birds were in the Search state in that square.

### Stopping might underestimate spatial memory

What role, then, do landmarks play in the spatial memory of these birds? If birds that had only a single training trial relied on the landmarks for almost all information about the flower’s location but acquired alternative cues for recognising the broad area in which the flower was situated as they learned about their surroundings, the landmarks would no longer be necessary for starting Search. The coarse information provided by the non-experimental cues, however, was no substitute for the fine-scaled information provided by the experimental landmarks 30cm from the flower. This explanation would account for the clear effect of the landmarks on the birds’ stopping locations. But stop locations would have to represent honest information about how well hummingbirds remember a location. The detailed analysis of the birds’ behaviour in Experiments 2 and 3, however, suggests that stops might actually underestimate spatial memory. For example, if we focus only on the *closest* distance that birds stopped from, and flew to, the flower’s location, we can see how landmarks affected the maximum accuracy the birds achieved. While the *closest stop* by birds with the landmarks was closer to the flower’s location than was the closest stop by the birds without the landmarks (z = 2.073, P = 0.038), these birds did not differ in the closest distance they *flew* to the flower’s location (z = 0.925, P = 0.36, Figure 2). The landmarks appear to be necessary for stopping close to the flower, but not necessary for flying close to the flower’s location.

If we then compare the difference between the closest stop and the closest distance flown from the flower, with landmarks present the two measures of a bird’s behaviour concur: the closest stop was usually within a few centimetres of the closest to the flower that a bird flew overall. But when the landmarks were removed this difference increased to tens of centimetres (z = 2.41, P = 0.016): stops and overall accuracy were decoupled. This decoupling did not require multiple training trials and was seen even after a single experience of the flower’s location (Experiments 1 & 2: t = - 5.563, P < 0.0001).

Determining “where to stop” and “where to search” might therefore rely on different kinds of information. The birds use the landmarks much more to inform where to stop than they use them to decide where to search, even though both form decisions are based on the hummingbird’s spatial memory of the flower’s location.

### General Discussion

When hummingbirds had a single experience of the flower’s location, birds heavily relied on familiar nearby landmarks both when choosing to initiate searching (Figure 1) and when choosing where to hover during searching (Figure 2). When birds had multiple experiences of the flower’s location, however, birds only relied on these landmarks when deciding where to hover and were able to use other cues when deciding where to start searching.

How hummingbirds use landmarks therefore not only depends on experience, but also on the kinds of decisions birds are making. Differences in the function of these decisions during navigation could explain changes in how reliant birds were on the experimental landmarks. Initiating search, for example, doesn’t need to be within centimetres of the goal, as birds only need to know they are close enough to slow down. Initially hummingbirds slowed down when they were a certain distance from the landmarks, but birds soon learned to use other cues to recognise when they were in a few metres of the goal. Stopping, on the other hand, could represent birds choosing to seek additional information, keeping the head and eyes still to clearly view the surroundings^13^ (similar to the “micro- choices” rats make before choosing a maze arm^14^). Ideally, birds would seek information closer to the goal, but the ability to do could depend on the resolution of the available spatial information. Without the landmarks, birds might be able to recognise when they are within a few metres of the goal based on the view of their surroundings, but the landmarks provide an unambiguous cue to the goal location even if only by making the view closer to the goal more distinctive^15,16^. It therefore makes sense that hummingbirds remained reliant on the landmarks for stopping but not for initiating search.

Studying decision-making in animal cognition must, therefore, involve more than just analysing the choices animals make. This development of landmark use was only discovered by looking beyond the easily quantifiable choices hummingbirds made (stopping) and looking at how movement as a whole changed as birds learned. Analyses of movements over much larger scales have long looked for “changepoints” which could reflect navigation decisions^17^, but these kinds of approaches are very rarely applied to smaller-scale experimental spatial memory tasks. Movement ecology tools, like HMMs, and advances in tracking technology provide an opportunity for comparative cognition to broaden its horizons, move beyond simple behavioural measures, and look closer at how animals acquire and use information across all stages of their behaviour.

## Supplementary materials

The material presented here is a supplement to “3D flightpaths reveal the development of spatial memory in wild hummingbirds” by David J Pritchard, T Andrew Hurly, Theoni Photopoulou and Susan D Healy (2021). **add doi**

### S1 Supplementary methods

#### S1.1 Subjects and experimental site

The subjects in this experiment were 14 male rufous hummingbirds. Each bird was individually identifiable by a coloured ink mark applied to his breast, and occupied a territory around one of 25 artificial feeders suspended 3m from the ground and positioned along the Westcastle valley of the Eastern Range of the Canadian Rockies in Alberta, Canada (49deg 29’N, 114deg25’W). This experiment was conducted between May and July 2014, when the males had migrated to Canada from Mexico for the breeding season. The experimental array consisted of a novel pink “test flower”, 3cm in diameter at the top of a 62cm long wooden pole, with two landmarks 30cm away placed with one to the northwest and one the northeast of the flower in an equilateral triangle. The landmarks were red cubes make of card and red duct tape, 10cm × 10cm × 10cm, on top of wooden poles 62cm long. The behaviour of hummingbirds was recorded using a pair of Sony Handicams, 1.5m apart and 5m away from the flower.

#### S1.2 Training and testing

##### S1.2.1 Experiment 1: single experience, landmarks present

Hummingbirds were trained to find 25% sucrose from a yellow “training flower”, 6cm in diameter, on the feeder. On the day each bird was to be tested, we removed the feeder and placed the training flower on a stake directly beneath the feeder’s location. We then positioned two Sony Handicam camcorders on tripods 1.5m apart and 5m from the training flower. Both cameras were centred on the location of the training flower. After the resident bird fed from the training flower, we started the cameras, removed the training flower, and set up the experimental array. The experimental array consisted of a novel pink “test flower”, 3cm in diameter at the top of a 62cm long wooden pole, with two landmarks 30cm away placed with one to the northwest and one the northeast of the flower in an equilateral triangle. The landmarks were red cubes make of card and red duct tape, 10cm × 10cm × 10cm, on top of wooden poles 62cm long. During the first test, the test flower was always placed 1m from the previous position of the yellow flower. We synchronised the cameras by hitting a red screwdriver against a white hammer. The point at which the screwdriver hit the hammer lasted less than 1/25th of a second, and so could be used to synchronise the videos during editing. We then calibrated the cameras by holding a 40cm × 40cm black and white chequerboard of 100 squares each one 4cm × 4cm, in 20-25 different locations visible to both cameras. we used the views of the chequerboard, as seen from each camera, to perform a stereo calibration of the cameras during analysis. During the first test, we removed the feeder from the hummingbird’s territory, and we allowed the territorial bird to visit feed once from the flower in the experimental array test flower only once. If the territorial bird did not feed from the test flower after multiple attempts, as was the case at three sites, we turned off the cameras, removed the test flower, allowed the hummingbird to feed 2-4 more times from the training flower at different locations more than 2m from the location of the experimental array. We then started the test again from the beginning with the training flower below the feeder. If, as at two sites, the birds continued to avoid the test flower, we trained the birds to visit the test flower. We did this by allowing the birds to feed from the training flower at a location more than 2m from the location of the experimental array. When the bird had fed once, we placed the pink disc of the test flower over the larger yellow disc of the training flower to make a hybrid flower, which we then allowed the birds to feed from once. After bird had fed from the hybrid flower, we removed the yellow disc so that only the test flower remained. After the bird had fed from the test flower, we moved the flower to a new location, again more than 2m from the location of the experimental array, and allowed the bird to feed again. Having fed from the test flower no more than four times, the flower was removed, the feeder replaced, and the test started again, from the beginning on the following day. After the bird had fed from the test flower and flown away, we removed the test flower, but kept the landmarks in place. We synchronised the cameras once more, before retreating to wait for the bird to return. Following the second visit by the bird to the experimental array with the flower removed, we synchronised the cameras one last time, removed the landmarks and replaced the feeder. Both visits were recorded by the cameras.

##### S1.2.2 Experiment 2: single experience, near or far, landmarks present or absent

After the bird had fed once from the feeder, we replaced the experimental array. For birds in the Near condition, we placed the test flower 40cm from the location of the test flower in the first test. For birds in the Far condition, we placed the test flower 80cm from the location of the test flower in the first test. As before, we placed the landmarks 30cm from the current location of the test flower, to the north in an equilateral triangle with the flower. We then started recorded, calibrated and synchronised the cameras. Once the territorial bird had fed once from the flower, we synchronised the cameras, removed the test flower, and, for birds in the “No Landmarks” group, we also removed the landmarks. For birds in the “Landmarks” group, the landmarks remained in place. After the bird had visited a second time and flown away, we synchronised the cameras for the last time, stopped recording, we removed the landmarks if they were present, and replaced the feeder. As in the first test, all visits were recorded by the cameras.

##### S1.2.3 Experiment 3: multiple experiences, landmarks present or absent

Once the territorial bird had fed once from the feeder, the experimental array was put back out in the same location as in second test. During the third test, we allowed the hummingbird to feed twelve more times from the test flower. Once the bird had fed and flown away after the twelfth visit, we synchronised the cameras, and removed the test flower. Following the eleventh visit, we started recording, calibrated and synchronised the cameras, and allowed the bird to feed for a twelfth time from the test flower. As in the second test, for birds in the “No Landmarks” group, we removed the landmarks alongside the flower, while for birds in the “Landmarks” group, the landmarks remained in place. After the bird had visited once more, we synchronised the cameras for a final time, we removed the landmarks if they were still out, stopped recording, and replaced the feeder. Due to battery constraints, in the third test we only recorded the twelfth visit and the test itself.

#### S1.3 Data extraction

We edited the videos in Sony Vegas, matching up the synchronisation points on the left and right videos, and splitting the videos in to three distinct sections: Calibration, training (the first visit, with the flower present), and testing (the second visit, with the flower removed). Having saved the new, synchronised video segments, we then calibrated the cameras and extracted the flight paths of the birds.

##### S1.3.1 Calibration

To reconstruct the location of the hummingbird, we stereo calibrated the cameras, determining their position, orientation, and skew. To perform the stereo calibration, we first used ffmpeg to extract the frames from each of the calibration videos, then used a custom MATLAB script to identify frames where the entire chequerboard was clearly visible and had been from three frames before to three frames after. Having identified good images of the chequerboard from both cameras, we used the Stereo Camera Calibration App in MATLAB to calibrate the cameras based on the view of the chequerboard from the left and right cameras in the chosen images. Because this method required the chequerboard to be rectangular, with an odd number of squares on one axis and an even number of squares on the other, one row of black squares was removed from the calibration images using a white brush in Microsoft Paint prior to calibration. The known dimensions of the chequerboard resulted in the absolute distances between the cameras, and by extension objects reconstructed after calibration, being accurate to the distance in real life.

##### S1.3.2 Flight path extraction

For each training and testing video segment, we extracted the frames from the videos, which we then sorted depending on what the bird was doing. In training videos, the frames were split into “In” (from first view of the bird to feeding from the flower), “Feeding”, and “Out” (from leaving the flower to disappearing from view). In test videos, we kept the entire period from the bird entering the shot, to the bird leaving. Using a custom MATLAB script, these images were converted to .avi video files, and imported into Kinovea to extract the trajectory of the birds from each of these videos by clicking on the head of the bird in each frame. When the bird was moving too fast to be seen clearly, we instead clicked on the leading edge of the blur seen as the bird moved. Overall, we collected six sets of trajectories for each test at each site, the head coordinates during the flight in as seen from each camera, the head coordinates during the flight out as seen from each camera, and the head coordinates during the fight in the test phase as seen from each camera. This resulted in a dataset of 2679 locations from 14 birds in experiment one, 1830 locations from 14 birds in experiment two, and 1604 locations from 13 birds in experiment three.

##### S1.3.3 Three-dimensional reconstruction

With the stereo calibration data from the cameras, and the x, y coordinates of the bird’s head from the videos, we were then able to reconstruct the x, y, z coordinates of the head of the bird in relation to the camera by using the triangulate function in the stereo calibration package.

#### S1.4 Statistical analysis of reconstructed trajectories

We used multivariate hidden Markov models (HMM) to analyse the reconstructed trajectories of the birds under each experimental setup. HMMs are models for analysing time series of observations recorded at regular time intervals. They are now commonly used in the animal movement context for inferring the underlying movement modes that give rise to observed movement metrics (e.g., [1]). The classical HMM formulation assumes a first-order dependence between the underlying states, and between the observations and the underlying states. HMMs for animal movement often involve modelling two derived variables (one scalar and one angular) from relocation data, e.g., the step length and turning angle between consecutive observed locations in the case of terrestrial movement. Flying animals move in a volume rather than on a plane, and this can easily be accommodated in HMMs by including an extra angular component, or data stream, as we do here. The trajectory reconstruction provided a three-dimensional step length, a yaw angle (nose side to side) and a pitch angle (nose up and down). We modelled step length using a gamma distribution, which is suitable for modelling continuous, positive-valued variables, as in step length. There were some steps which had zero length (no displacement) and this was accounted for by estimating an additional parameter for the probability mass of the observations that are zero. We modelled both angular state variables using a wrapped Cauchy distribution, which has two parameters, the mean and concentration, where the latter relates to how peaked the distribution is. We fitted models where either both parameters were estimated, or only the concentration parameter while assuming the mean to be zero.

##### S1.4.1 Model structure and implementation

An unobserved Markov chain was assumed to determine the behavioural states and the parameters of the state-dependent distributions associated with the observed movement variables. Our research question pertains to inferring information about the different types of movement behaviour carried out by rufous hummingbirds. We fit the model to all birds jointly and therefore estimate one set of parameters across individuals. We also fitted the model found to perform best for all birds for a given experiment, to each bird separately for that experiment. The HMM parameters are estimated using numerical maximisation of the likelihood, implemented in R [2], using the momentuHMM package [3]. In this package, the computation of the covariate-dependent transition probability matrices and the forward algorithm are coded in C++. The forward algorithm is an efficient way of evaluating the likelihood and is one reason for the popularity of HMMs – it makes them fast to fit. It corresponds to a recursive calculation of the likelihood with computational costs only linear in the number of observed time points and renders numerical maximum likelihood estimation feasible [4]. We only fit models with two states, based on biological knowledge of the movement behaviour of the birds during the experiment, and guided by the aim of the analysis. We expected birds to exhibit more rapid, directed movement towards an area of interest, followed by more sinuous and perhaps slower movement while searching for a flower to feed from. We did not consider models with different numbers of states, following the pragmatic approach outlined in [5].

##### S1.4.2 Likelihood of the HMMs

The likelihood of an HMM with N states and observation vectors, …, can be written as a matrix product:

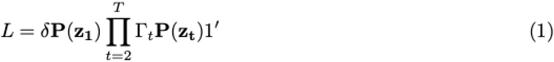

where δ is a row-vector containing the initial state distribution, represents the N×N transition probability matrix at time point t and 1 is a row-vector of ones. P denotes a N ×N diagonal matrix containing the values of the N joint state-dependent densities evaluated at the observation vector. We assume the observed variables to be contemporaneously conditionally independent, given the current state. Thus, for each state, the joint state-dependent density is the product of the univariate state-dependent densities which are associated to the observed variables. In this analysis, corresponds to the vector: step length, yaw angle (side-to-side movement), and pitch angle (up-down movement) observed at time t along the movement track. We model the three state variables (step, yaw, pitch) with distance between the bird and the test flower (referred to as current distance to flower from here on) as a covariate on the probability of transitioning between states. We also included the presence of landmarks as a covariate in experiments two and three - landmarks were present for all birds in experiment one. We implemented this in two ways, 1) we included the presence of landmarks as a covariate on the state dependent distributions, to test whether the parameters of the estimated component distributions of movement states change when landmarks are present, and 2) we included the presence of landmarks as a covariate on the transition probabilities, the same as with current distance to flower. To investigate the influence of distance between the bird and the previous location of the flower (m), and the effect of the presence of landmarks (0/1) on movement behaviour, these two covariates are incorporated into the model either on the transition probabilities (independently or as interacting covariates) (Equation 2) or on the mean of the state-dependent distribution for step length (Equation 3).

For a model with distance from flower and the presence of landmarks interacting to affect the transition probabilities, the linear predictor would be

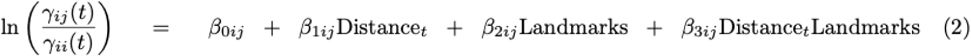

where denotes the probability of switching from state i to state j at time t. The effect of absence of landmarks is absorbed into the intercept term. For a model with the presence of landmarks acting on the mean of the state-dependent distribution for step length, the linear predictor would be

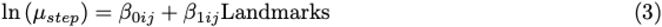

where, again, the absence of landmarks is absorbed into the intercept, and the slope coefficient corresponds to the presence of landmarks. Given we are not fitting any random effects, the log-likelihood of interest is the sum of log-likelihoods corresponding to the different birds within each experiment. S1.4.3 Model assessment

We used the AIC weights to choose between models with different covariates for each of the three experiments, but all models had two states. For the best model from each experiment, we used the Viterbi algorithm to obtain the most likely state sequence and we also calculated the state probabilities, which gives the probability of being in a given state at each point in the observed time series. We examined model fit for each experiment by calculating the pseudo-residuals for each of the state-dependent variable (step length, yaw, pitch) and checking their distributions and the residual autocorrelation, using the acf function in R [2].

##### S1.4.3 Analysis of stops and distances

For the generalised linear mixed models, the response variable was standardised, scaled and a constant added to avoid negative values. We fitted all possible combinations of explanatory variables using the dredge function in MuMIn, which compares models based on AICc, delta weight, and Akaike weight. In cases in which the delta weight of between models was less than 2, model outputs were averaged using the ‘model.avg’ function in the MuMIn package.

### S2 Supplementary results

#### S2.1 Candidate models

In the main text we presented only the top few models for each experiment (Table 1, main text). Here we describe all models that we tried for each experiment, including their AIC scores and AIC weights (Table S1).

#### S2.2 Diagnostics on best models

The distributions of the pseudo-residuals for the two angular state-dependent variables (yaw, pitch) were symmetrical and showed no evidence of residual autocorrelation. The pseudo-residuals for step length, especially in experiment three, deviated from symmetry, suggesting that there are perhaps additional movement states that become more pronounced when spatial memory is a strong component motivating movement, for example, stopping behaviour. We made a conscious choice not to fit models with more than 2 states for biological interpretability in the face of a relatively small dataset for each experiment. Step length showed some residual autocorrelation in all three experiments but dropped down to within the confidence bounds of zero within 10 time steps.

#### S2.3 Individual variability

To explore individual variability we took the best fitting model, based on the data from all birds combined for each experiment, and fitted it to each bird individually. The results are presented in Figure S1. In experiment 1, all birds had the landmarks present and the best-fitting model included distance of the bird from the previous location of the flower as a covariate on the transition probability matrix. We did not consider including landmarks as a covariate in the model for this experiment because they were present for all birds. In experiment 2, some birds had landmarks present and some had landmarks removed. The best-fitting model had the interaction of distance from flower and the presence of landmarks as interacting covariates on the transition probability matrix. In experiment 3, some birds had landmarks present and some had landmarks removed but the best-fitting model did not include the presence of landmarks as a covariate on the transition probability matrix so it has no effect on the transitions shown in Figure S1. The individual models did not converge for birds 3 and 10 in the no landmarks treatment in experiment 2. This is likely due to the small number of locations from each of these two birds, 45 and 24 respectively. The model for bird 2 from experiment 3 also did not converge for unknown reasons.

